# Comparing manual and automatic artifact detection in sleep EEG recordings

**DOI:** 10.1101/2023.05.14.540709

**Authors:** Ujma P. Péter, Martin Dresler, Róbert Bódizs

## Abstract

All sleep EEG recordings can be contaminated by artifacts. Both visual and automatic methods have been developed to mark such erroneous segments of EEG data. Here we systematically explore the effect of artifacts on the sleep EEG power spectrum density (PSD), and we compare gold-standard visual detections to a simple automatic detector using Hjorth parameters to identify artifacts. We find that most distortions in the all-night average PSD occur because of a small minority of highly anomalous artifacts, which mainly affect the beta and gamma frequency ranges and NREM delta. Visual and automatic detections only show moderate agreement in which data segments are artefactual. However, the resulting all-night average PSD is highly similar across all methods, and PSDs calculated with all methods successfully recover the known correlations of PSD with age and sex. No parameter settings of the automatic detector clearly outperformed others. Additionally, we show that accurate average PSD estimates can be recovered from just a fraction of available data epochs. Our results suggest that artifacts represent a minor and easily solvable problem in sleep EEG recordings. Most visually identified artifacts do not seriously distort estimates of mid-frequency activity in the sleep EEG spectrum, and distortions to low and high frequencies can be eliminated using a simple automatic detection method nearly as well as with visual detections. These findings show that the visual inspection of EEG data is not necessary to eliminate the effects of artifacts, which is encouraging for the expected performance of automatic preprocessing in large sleep EEG databases.

## Introduction

With the advent of large, freely accessible databases, there is a renewed interest in the sleep electroencephalogram (EEG) as a source of disease biomarkers and features related to normal brain function (Zhang et al., 2018; Redline and Purcell, 2021). Sleep EEG data is free from blinking artifacts and less contaminated by movement artifacts than wakeful EEG, however, various artifacts related to muscle activity during arousals, cardiac activity, the temporary disconnection of electrodes or perspiration may contaminate it (Islam et al., 2016). In small sleep EEG studies it is possible to visually detect artifacts, that is, have a person screen all EEG data and mark up artefactual epochs by hand. However, this task is increasingly impractical in case of large databases (Mariani et al., 2018). Various automatic detection methods have been developed to detect artifacts in sleep EEG data, using various features such as the slow/fast power ratio (Mariani et al., 2018), abnormal amplitude or power (Brunner et al., 1996; D’Rozario et al., 2015) or Riemannian geometry (Saifutdinova et al., 2019) (see (Anderer et al., 1999; Motamedi-Fakhr et al., 2014; Urigüen and Garcia-Zapirain, 2015; Islam et al., 2016; Malafeev et al., 2019; Cox and Fell, 2020) for reviews).

Many previously published automatic artifact detection algorithms achieved high agreement with visual detections (t Wallant et al., 2016; Mariani et al., 2018; Malafeev et al., 2019; Saifutdinova et al., 2019), https://doi.org/10.1111/jsr.12679and were successful in developing automatic detection algorithms which resulted in a significant improvement in signal quality. However, to our knowledge there is no empirical study which systematically investigated how featured calculated from the sleep EEG are affected by the erroneous inclusion of artefactual data (but see (Malafeev et al., 2019) for a partial analysis and (Brunner et al., 1996) for the systematic description of the effect of muscle artifacts), and there is no consensus about simple and effective automatic methods which can replace the visual scorings of recordings before analysis, even though this would be critical in large datasets where visual scoring is not feasible. In the current study, especially designed to bridge this gap, we use a large sample of healthy volunteers to compare visual detections with a simple automatic method pioneered in recent analyses of large sleep EEG datasets available at sleepdata.org (Purcell et al., 2017; Djonlagic et al., 2021). We show that a minority of visually scored artifacts are responsible for most of the distortions of EEG spectra, these are readily captured by a simple method based on finding epochs with outlying Hjorth parameters, and the resulting PSD estimates are similar to those obtained with visual scorings and reliably demonstrate similar correlations with age and sex., and were successful in developing automatic detection algorithms which resulted in a significant improvement in signal quality. However, to our knowledge there is no empirical study which systematically investigated how featured calculated from the sleep EEG are affected by the erroneous inclusion of artefactual data (but see (Malafeev et al., 2019) for a partial analysis and (Brunner et al., 1996) for the systematic description of the effect of muscle artifacts), and there is no consensus about simple and effective automatic methods which can replace the visual scorings of recordings before analysis, even though this would be critical in large datasets where visual scoring is not feasible. In the current study, especially designed to bridge this gap, we use a large sample of healthy volunteers to compare visual detections with a simple automatic method pioneered in recent analyses of large sleep EEG datasets available at sleepdata.org (Purcell et al., 2017; Djonlagic et al., 2021). We show that a minority of visually scored artifacts are responsible for most of the distortions of EEG spectra, these are readily captured by a simple method based on finding epochs with outlying Hjorth parameters, and the resulting PSD estimates are similar to those obtained with visual scorings and reliably demonstrate similar correlations with age and sex.

## Methods

### Participants and EEG acquisition

We used all-night EEG data from 252 healthy volunteers (mean age=25.14 [SD=12.21, range 3.8-69], 130 males, 122 females). The technical details of EEG acquisition in this dataset were previously reported (Bódizs et al., 2017, 2022). In short, this is a multicenter dataset of the Max Planck Institute of Psychiatry (Munich, Germany) and the Psychophysiology and Chronobiology Research Group of Semmelweis University (Budapest, Hungary). Participants underwent all-night polysomnography recordings for two consecutive nights. We used data from the second night, from EEG channels common to all recordings: Fp1, Fp2, F3, F4, C3, C4, P3, P4, O1 and O2, all referenced to the mathematically linked mastoids. Hypnograms for all recordings were visually scored on a 20 second basis according to standard criteria (Iber et al., 2007). In the current version of the dataset, an additional 28-year-old male was included in the “MPIP-I” subsample.

Study procedures were approved by the ethical boards of Semmelweis University, the Medical Faculty of the Ludwig Maximilian University or the Budapest University of Technology and Economics. All participants were volunteers who gave informed consent in line with the Declaration of Helsinki. In case of participants who were minors, written informed consent was provided by the parents.

### Artifact detection and power spectral density (PSD) analysis

Manual artifact detection was completed in all recordings based on visual inspection of the EEG waveforms by experts in sleep EEG scoring and analysis. This detection was performed on a 4-second basis. Each 4-second epoch was either confirmed as good quality or rejected, with this scoring applied to all channels. Artifact scoring was performed to exclude all visual identifiable artifacts, including physiological/biological (myogenic, eye or body movement-related, teeth grinding, sweating, respiration or pulse-related) and technical ones (electrode pop) were excluded from quantitative EEG processing.

Automatic artifact detection was based on Hjorth parameters (Hjorth, 1970), used for artifact detection in several recent studies of large-scale sleep datasets (Purcell et al., 2017; Djonlagic et al., 2021). Hjorth parameters are computationally simple indicators of the statistical properties of time series signals. The three Hjorth parameters are activity, mobility and complexity.

Activity is defined as

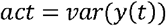

where y(t) is the signal. In other words, activity is simply the variance of the signal amplitude, which according to Parseval’s theorem equals the total area under the power spectral density curve.

Mobility is defined as

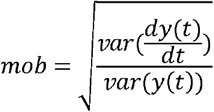

, where y(t) is the signal and dy(t)/dt is the first derivative of the signal. Mobility reflects the average slope of the EEG relative to overall signal amplitude (preponderence of sudden changes in EEG amplitudes lead to higher mobility).

Complexity is defined as

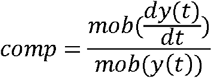

In other words, complexity is the mobility of the first derivative of the signal divided by the mobility of the signal. The minimal value (comp = 1) of this ratio would emerge in the case of a pure sine wave (no complexity).

We calculated all three Hjorth parameters for all 4-second epochs in each participant, on each channel, separately in NREM and REM using custom code written in MATLAB R2021a. The code used in analyses and visualization is available https://zenodo.org/record/7934586. Raw data is also accessible using the resources posted here.

Hjorth parameters follow a continuous distribution, but the decision whether to include an epoch in analyses is binary. We explored how individual average spectra calculated using various decision criteria relate to the gold standard visual detection method. First, we calculated within-participant, within-channel z-scores of all Hjorth parameters across all 4-second epoch within the same vigilance state (NREM or REM). Second, we calculated individual mean spectra only considering epochs in which z-scores did not reach a z-score threshold. These thresholds were set at z=1, 1.5, 2, 2.5, 3 and 4, representing six increasingly lax criteria for which epochs are considered artefactual. In a further step, we explored a dual-threshold method where z-scores were re-calculated after artefactual epochs were rejected by the initial threshold, and some of the remaining epochs were rejected again based on a new threshold applied to the updated z-scores (Purcell et al., 2017). The rationale behind this approach is that the first threshold removes gross outliers and the second targets more moderate artifacts. For the dual-threshold approach, we used five thresholds of z=1/1, 1.5/1.5, 2/2, 2.5/2.5 and 3/3 for the first and second thresholds, respectively. In total, 13 NREM and REM individual average power spectra were calculated for each participant and each channel, one with visual artifact rejection, one with no artifact detection at all, and eleven with different Hjorth parameter thresholds applied in the automatic analyses. Power spectra were log10 transformed, but not relativized. Where analyses were performed within frequency bands, we used the following band criteria: delta (0.5-4 Hz), theta (4-8 Hz), alpha (8-10 Hz), low sigma (10-12.5 Hz), high sigma (12.5-16 Hz), beta (16-30 Hz), gamma (30-48 Hz). Frequency bins at the boundaries were always included in the lower frequency band.

Power spectral density was calculated for each 4-second epoch in the data (with 2 second overlap) using the periodogram() MATLAB function with a Hamming window. Average power spectra of the night were calculated by calculating the average of all non-artefactual epochs. An epoch was considered as non-artefactual (in both visual and automatic detections) if neither its starting point nor its end point was included in an epoch marked as artefactual. Average power spectra were log-transformed for analyses.

### Statistical analysis

In our analyses, we explored 1) the agreement between visual and automatic artifact detection methods on the epoch level, 2) the similarity of the individual average PSD across visual and automatic methods, 3) the extent to which the correlations of individual average PSD with age and sex can be recovered using various detection methods. Visual and automatic artifact detections were compared using Cohen’s kappa. PSD similarity was calculated by calculating the Pearson correlation (between participants) of the log10-transformed PSD estimates resulting from visual and automatic detections in each frequency bin. Pearson correlations between log10-transformed PSD estimates and age and sex were also calculated for each frequency bin. Sex was binary-coded as 1 for males and 0 for females, rendering the calculated correlation to be a point biserial correlation, with negative coefficients suggesting higher PSD estimates in males.

## Results

### Distribution of visually identified artifact effects

To simulate how much it affects average PSD estimates if a visual or automatic artifact detector misses errors in the data, we calculated the average PSD with each visually identified artifact erroneously included in analyses (but leaving other artifacts excluded). The result of these simulations is illustrated on **Figure 1** in NREM and **Figure 2** in REM.

**Figure 1.**
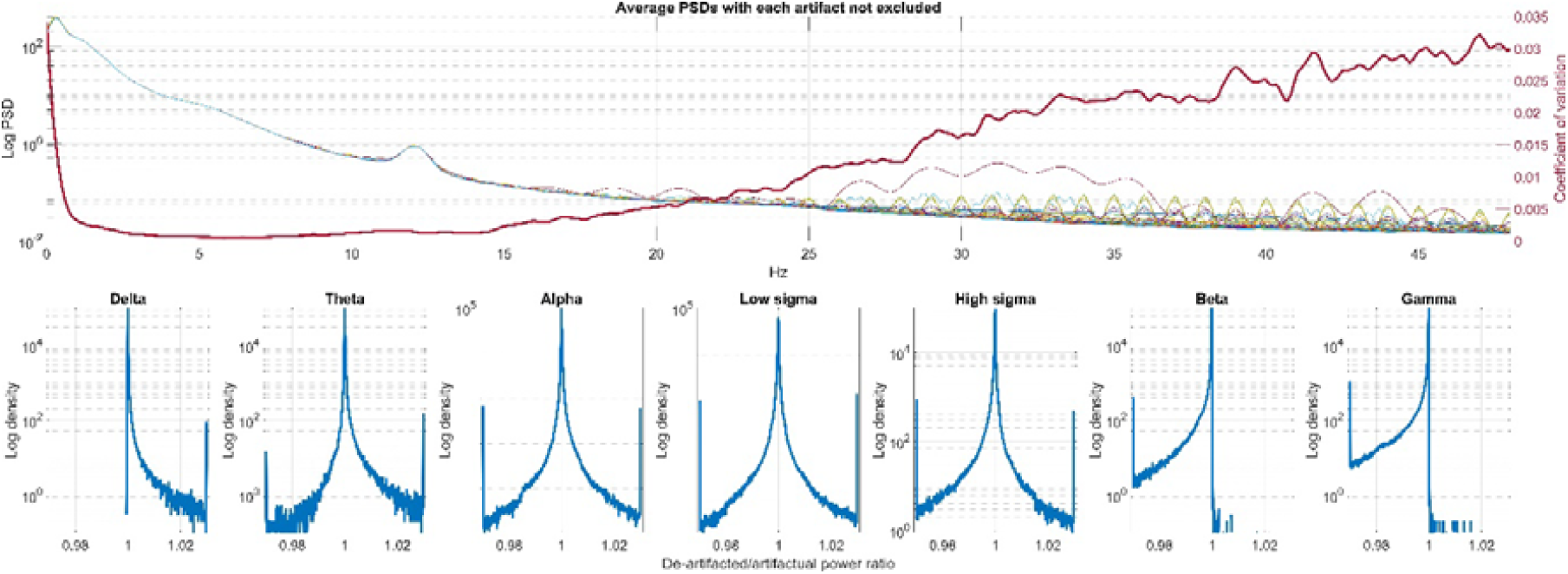
The effect of artifact non-exclusion to NREM PSD estimates. **Panel A** shows the average PSD estimate from a random participant and a random channel (O2) if one of the artifacts (but not the others) are erroneously included. Note that the lines almost perfectly overlap, except for higher-frequency activity. The coefficient of variation (standard deviation of artifact estimates divided by the mean, calculated separately for each frequency bin) gives a quantitative estimate of this effect. Coefficient of variation was calculated from each participant from the same channel and averaged across participants. **Panel B** shows histograms of the biasing effects of artifacts, estimated as the de-artifacted/artefactual log PSD summed across all bins in the frequency range. Ratios are shown by frequency range and pooled across participants and channels (Between-channel variance was minimal and adding between-channel SD shading was not visible). Ratios <0.97 and >1.03 – over 3% change in the final PSD estimate with the inclusion of an artifact – were set to 0.97 and 1.03, respectively. Note the symmetrical distribution of the mid-frequency artifact effects, but highly asymmetrical delta and beta/gamma effects.

**Figure 2.**
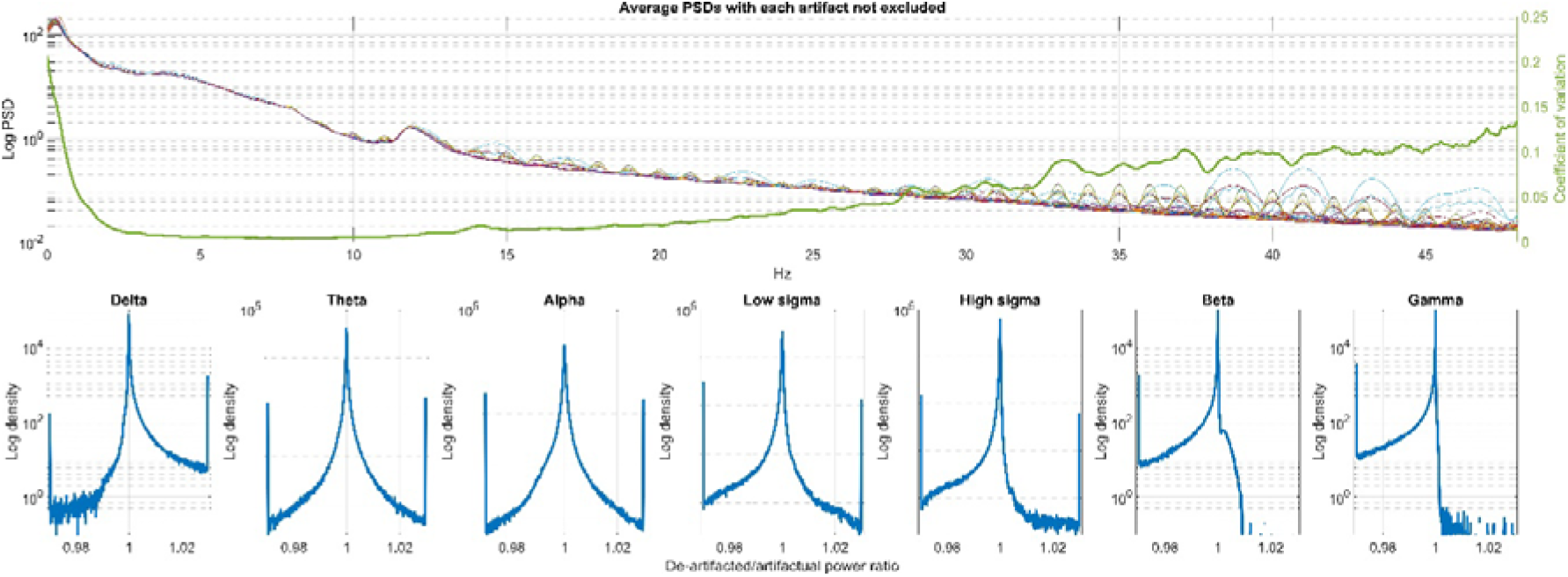
The effect of artifact non-exclusion to REM PSD estimates. **Panel A** shows the average PSD estimate from a random participant and a random channel (O2) if one of the artifacts (but not the others) are erroneously included. Note that the lines almost perfectly overlap, except for higher-frequency activity. The coefficient of variation (standard deviation of artifact estimates divided by the mean, calculated separately for each frequency bin) gives a quantitative estimate of this effect. Coefficient of variation was calculated from each participant from the same channel and averaged across participants. **Panel B** shows histograms of the biasing effects of artifacts, estimated as the de-artifacted/artefactual log PSD summed across all bins in the frequency range. Ratios are shown by frequency range and pooled across participants and channels (Between-channel variance was minimal and adding between-channel SD shading was not visible). Ratios <0.97 and >1.03 – over 3% change in the final PSD estimate with the inclusion of an artifact – were set to 0.97 and 1.03, respectively. Note the symmetrical distribution of the mid-frequency artifact effects, but highly asymmetrical delta and beta/gamma effects.

In both vigilance states, missing individual artifacts made little difference to the final PSD estimate in the ∼2-15 Hz frequency range. The coefficient of variation – indicating how much the final PSD estimate varies after the erroneous inclusion of individual artifacts – started notably increasing over 30 Hz in REM, but already at over 15 Hz in NREM, with an additional increase in the low frequency ranges. The modal artifact made minimal difference to the PSD estimates if erroneously included in analyses. However, a significant minority affected PSD estimates by more than 3% - note that this is the effect of misidentifying a single artifact out of thousands of epochs. In the theta, alpha and sigma ranges and in REM delta, the effect of artifacts on PSD estimates was symmetrical, with some artifacts causing an under- and others and overestimate of PSD. This was, however, not the case for NREM delta and for beta and gamma in both vigilance states. NREM delta PSD estimates were almost always lower and NREM/REM beta and gamma PSD estimates almost always higher if artifacts were erroneously included. In other words, artifacts in NREM contained less low-frequency activity than normal data, while in both vigilance states artifacts contained more high-frequency activity.

### Visual and automatic artifact detections show moderate agreement

Various metrics of detection agreement are shown in **Table 1**.

**Table 1.** Artifact detection characteristics. This table shows the proportion of epochs marked as artifacts in each method (“artifact rate”), agreement with visual detections based on Cohen’s kappa (“Detection vs. visual”), and the PSD correlations (“PSD vs. visual”) between the visual detection method and each alternative. Cohen’s kappa values are averaged across channels. PSD correlations are averaged across channels and frequency bins. For automatic detections, Hjorth parameter thresholds are expressed in standard deviation units. In case of dual thresholds, the two thresholds are separated by /.

|  |  |  | Hjorth parameter thresholds |  |  |  |  |  |  |  |  |  |  |  |
| --- | --- | --- | --- | --- | --- | --- | --- | --- | --- | --- | --- | --- | --- | --- |
|  |  | Visual detection | 1 | 1.5 | 2 | 2.5 | 3 | 4 | 1 / 1 | 1.5 / 1.5 | 2 / 2 | 2.5 / 2.5 | 3 / 3 | No artifact detection |
| NREM | Artifact rate | 0,08 | 0,31 | 0,13 | 0,06 | 0,03 | 0,02 | 0,01 | 0,72 | 0,32 | 0,15 | 0,09 | 0,05 |  |
|  | Detection vs. visual |  | 0,09 | 0,18 | 0,22 | 0,22 | 0,20 | 0,16 | 0,03 | 0,11 | 0,19 | 0,22 | 0,22 |  |
|  | PSD vs. visual |  | 0,97 | 0,96 | 0,95 | 0,94 | 0,93 | 0,90 | 0,96 | 0,97 | 0,97 | 0,97 | 0,96 | 0,78 |
| REM | Artifact rate | 0,18 | 0,23 | 0,14 | 0,09 | 0,05 | 0,03 | 0,01 | 0,58 | 0,29 | 0,19 | 0,14 | 0,09 |  |
|  | Detection vs. visual |  | 0,15 | 0,20 | 0,20 | 0,18 | 0,15 | 0,11 | 0,06 | 0,16 | 0,23 | 0,24 | 0,22 |  |
|  | PSD vs.<br>visual |  | 0,96 | 0,93 | 0,91 | 0,88 | 0,85 | 0,81 | 0,97 | 0,97 | 0,97 | 0,95 | 0,93 | 0,67 |

Visual artifact detection marked on average (across participants and channels) 8% of epochs in NREM and 18% of epochs in REM as artefactual. This rate was overestimated by the least stringent and generally underestimated by the most stringent automatic criteria. The best-performing automatic detector was the 2.5 SD dual-threshold method in both NREM in REM.

We calculated Cohen’s kappa between the binary artifact vectors produced by the visual method and those produced by automatic methods. This measure of agreement remained low regardless of the method used. The highest agreement in NREM (0.22) achieved by several threshold settings, and in REM the z=2.5/2.5 dual threshold method produced the highest agreement (0.24).

In order to estimate what recording characteristics affected the similarity of visual and automatic detection, we fitted a mixed-effects linear model using the fitlme() function in MATLAB, with Cohen’s kappa values between automatic and visual detections as the dependent variable, participant age, recording channel, and artifact detection method as fixed effects, and a random intercept per participant. Results are detailed in Table 2. Briefly, we found that lower participant age, NREM vigilance state and a more posterior channel location, were associated with lower agreement between visual and automatic detections. Less stringent detection criteria, especially with dual thresholds, were also associated with higher agreement. Effect sizes, however, were generally very low.

**Table 2.**
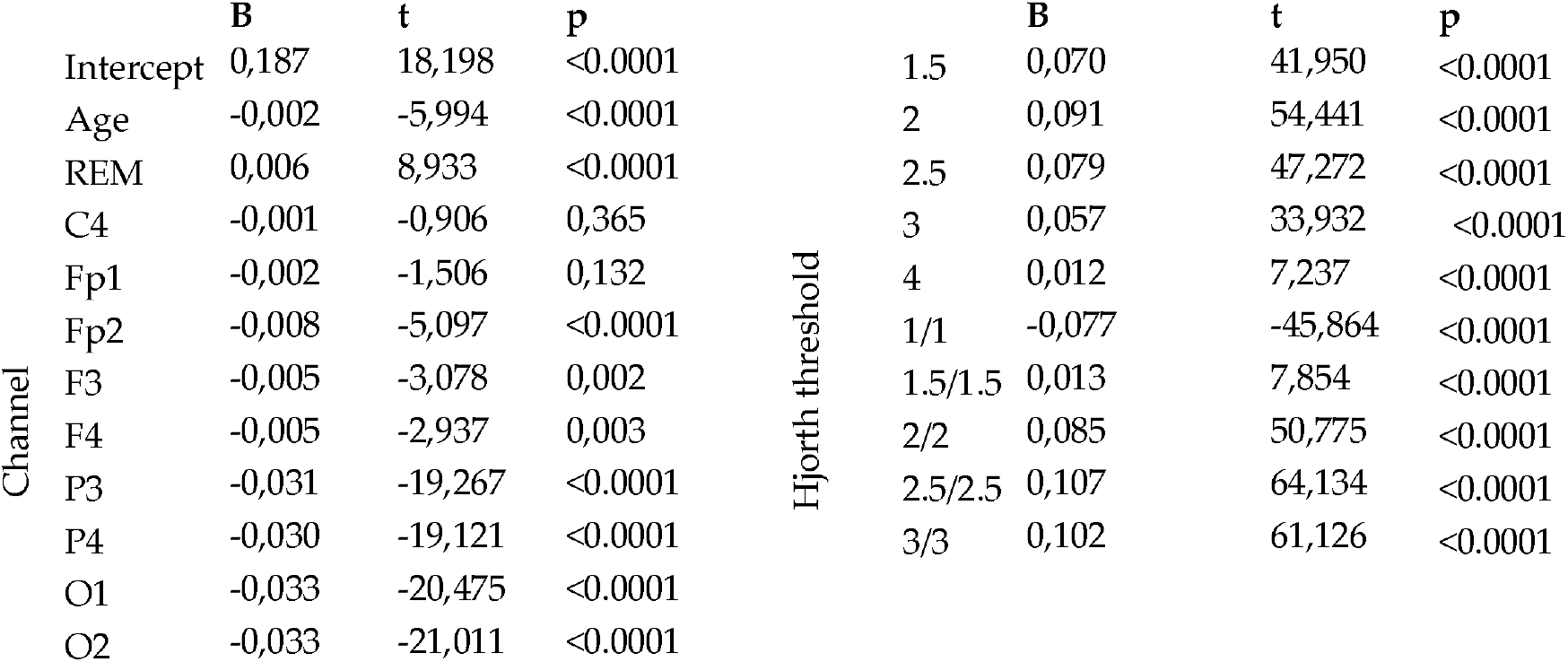
The effect of patient age, vigilance state, recording channel and the Hjorth parameter threshold used in automatic analyses on the agreement between visual and automatic detectors. Categorical variable effects are expressed relative to the reference categories NREM, channel C3 and Hjorth parameter threshold z=1.

Raincloud plots (Allen et al., 2019) illustrating the similarity of automatic detections with various thresholds on various channels to the visual detection gold standard are available in the Supplementary Material for NREM **(Supplementary Figure S1)** and REM **(Supplementary Figure S2)**.

### Different methods result in similar average power spectra

In the next step, we explored the similarity between average PSDs calculated using visual and automatic artifact detection methods. For this, we extracted the PSD estimate at each 0.25 bin between 0-48 Hz from each participant produced by the two methods (visual method and the automatic method to be compared with it) and calculated Pearson correlations between the two estimates. **Figure 3** shows the resulting estimates in NREM and **Figure 4** in REM. (See also **Table 1** for more coarse estimates).

**Figure 3.**
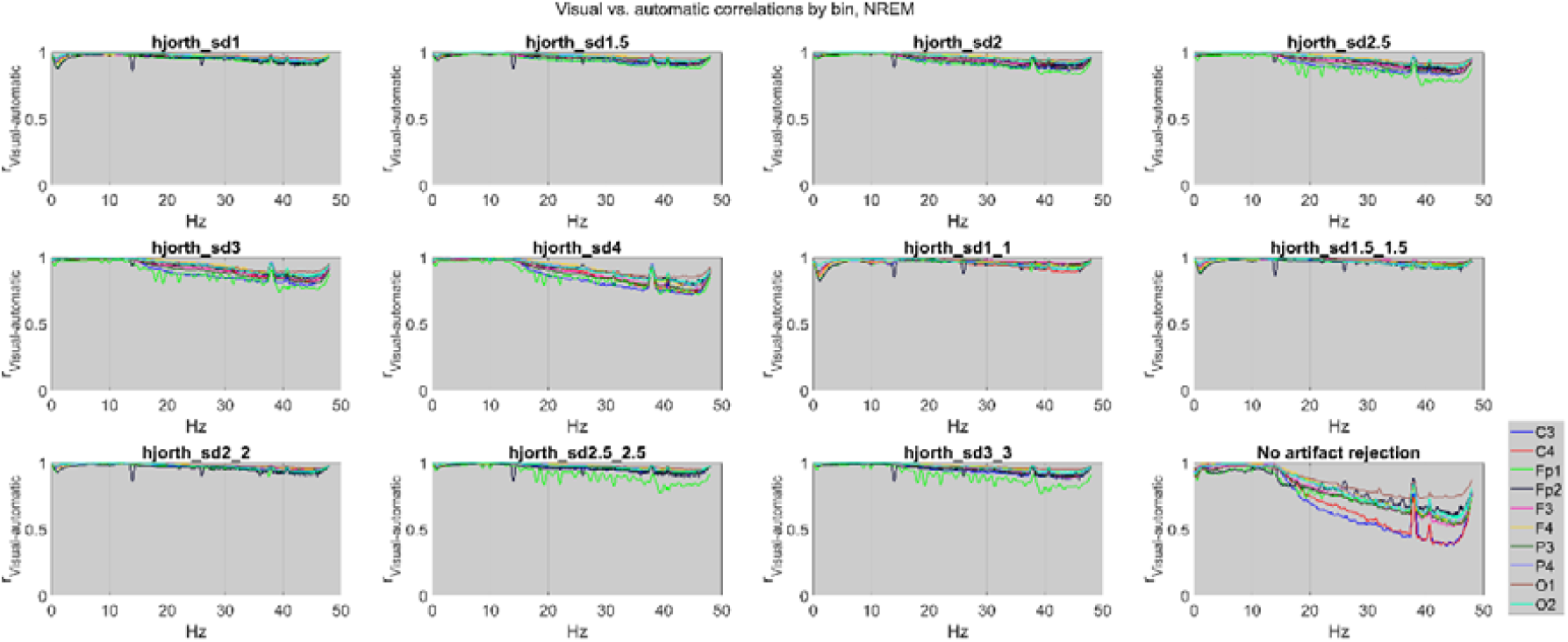
The similarity of binwise NREM PSD estimates obtained using visual and automatic detections. Line plots illustrate the Pearson correlation coefficient (between participants) between the PSD estimates obtained with visual detections and each automatic detector. Correlations are shown for each frequency bin and EEG channel. The last panel illustrates the similarity of PSD estimates with visual detections to a case where all epochs are included in analyses without regard for artifacts.

**Figure 4.**
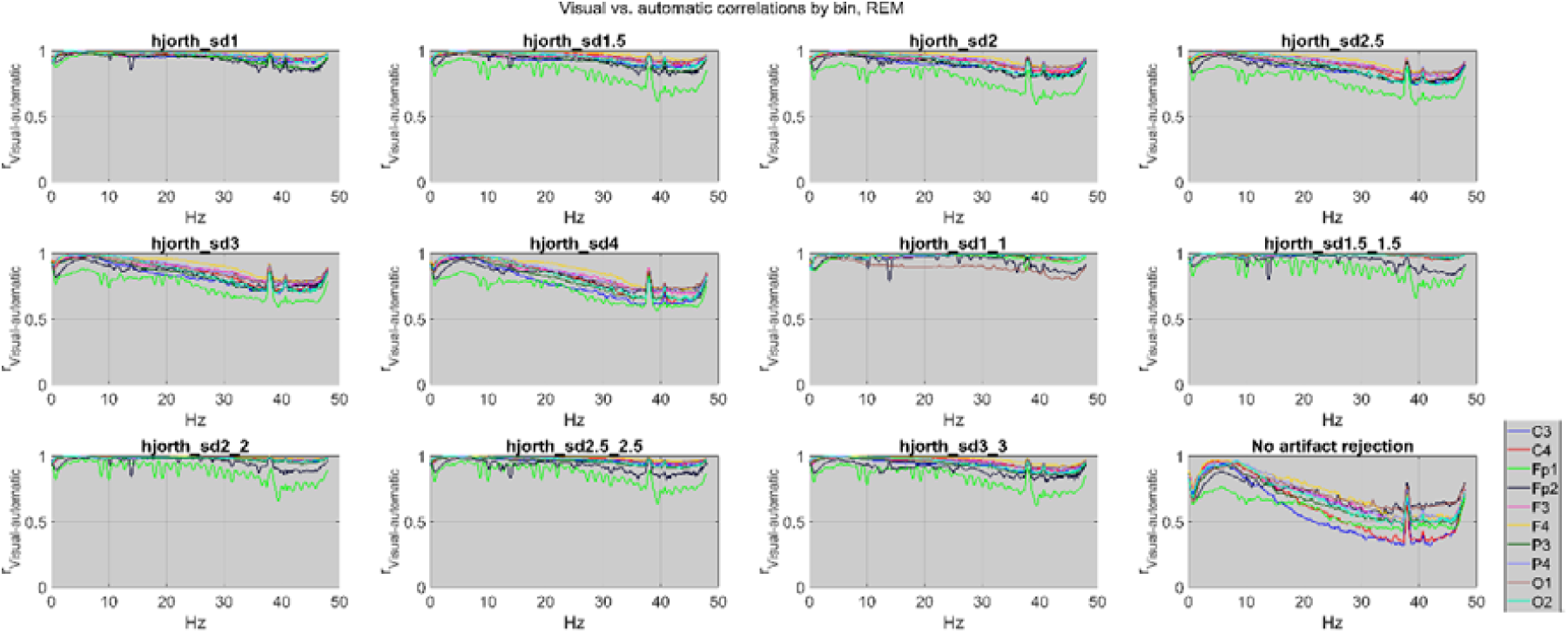
The similarity of binwise REM PSD estimates obtained using visual and automatic detections. Line plots illustrate the Pearson correlation coefficient (between participants) between the PSD estimates obtained with visual detections and each automatic detector. Correlations are shown for each frequency bin and EEG channel. The last panel illustrates the similarity of PSD estimates with visual detections to a case where all epochs are included in analyses without regard for artifacts.

PSD correlations between the visual and automatic were very high at least in the high delta-sigma range, especially in NREM, almost always exceeding 0.9 and often 0.95. Notably, very high correlations were obtained in this range even if no artifact detection method was used at all. Visual-automatic correlations were lower in the REM and above the sigma range, especially if no artifact detection method was used. While dual-threshold methods did not show better agreement with visual detections than single-threshold methods (see previous section), they did produce noticeably higher mean PSD correlations, indicating that they keep epochs which are more representative of those a visual observer would keep.

### External correlations

In subsequent analyses, we explored to what extent known correlates of individual average PSD can be reproduced using various artifact detection methods. We calculated correlations between PSD and age and sex.

Using the gold standard visual detection method, known correlations (Muehlroth and Werkle-Bergner, 2020), including those reported in this dataset (Pótári et al., 2017; Ujma et al., 2022) emerged. Higher age was associated with less low-frequency and spindle activity in NREM, and a reduction of low and an increase in beta-frequency activity in REM. Men had lower power across all frequency ranges in both NREM and REM, except the lowest and highest frequencies.

In line with the high correlations seen for PSD estimates, these patterns were generally well-recovered using various artifact detection methods, including no artifact detection at all. As expected, correlations with high-frequency components – such as age-related increases – were less well reproduced without artifact detections, but clearly seen once any automatic method was applied. External correlations are illustrated on **Figure 5**. More detailed plots, showing results for each channel separately, are available in **Supplementary Figure S3** (NREM, age), **Supplementary Figure S4** (NREM, sex), **Supplementary Figure S5** (REM, age) and for REM on **Supplementary Figure S6** (REM, sex).

**Figure 5.**
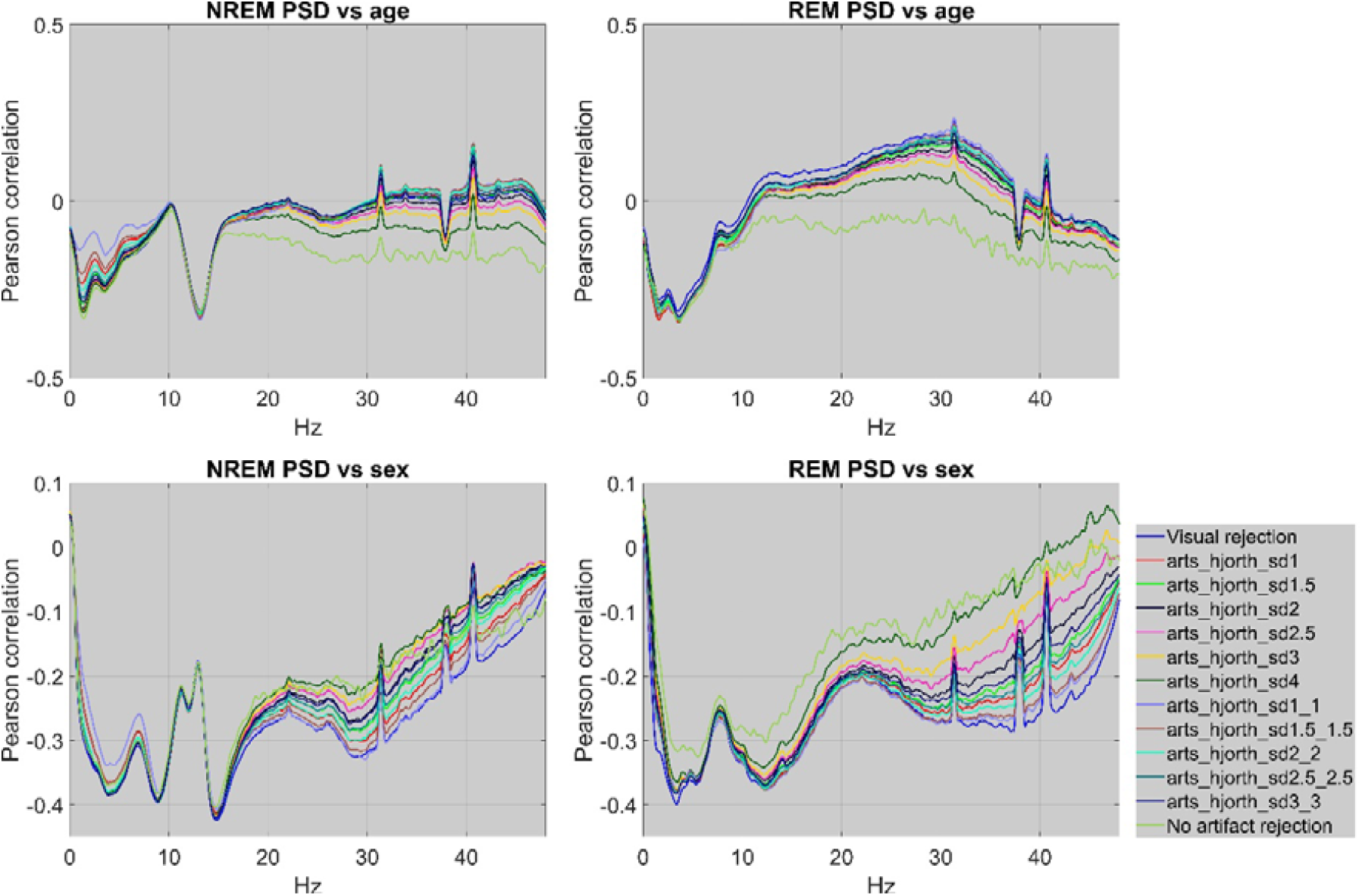
Correlations of PSD estimates with external indicators. The line plots show Pearson correlation coefficients between age, sex, and PSD estimates obtained using different artifact detection methods (including ignoring artifacts completely). Males are coded as 1 and females as 0, with a negative correlation implying higher PSD in females. Correlations are averaged across channels. Legend entries for automatic methods refer to using one (single number) or two (double number) SD thresholds to identify artifacts using outlying Hjorth parameters. Note the high overlap between lines, especially at the lower frequencies, indicating that the correlation of PSD estimates to external indicators is well-recovered regardless of the artifact detection method used.

### Simulating poor quality recordings

In our final analyses, we investigated the effect of excess artifacts on PSD estimates. For these analyses, we compared original PSD estimates to simulations where we randomly excluded epochs, emulating poor quality recordings or false positive artifacts detections by a visual scorer or an automatic detector.

In each participant, we set a random selection of epochs constituting 25%, 50% or 75% of good-quality data to artefactual. False positive artifacts were either selected from anywhere in the night or the first and second halves, respectively. We calculated PSD estimates using these artificial epoch sets and compared them with actual PSD estimates. Bandwise PSD averages were calculated by averaging frequency bins belonging to the same frequency band.

Results from the representative channel C3 are summarized on **Figure 6** for NREM and **Figure 7** for REM.

**Figure 6.**
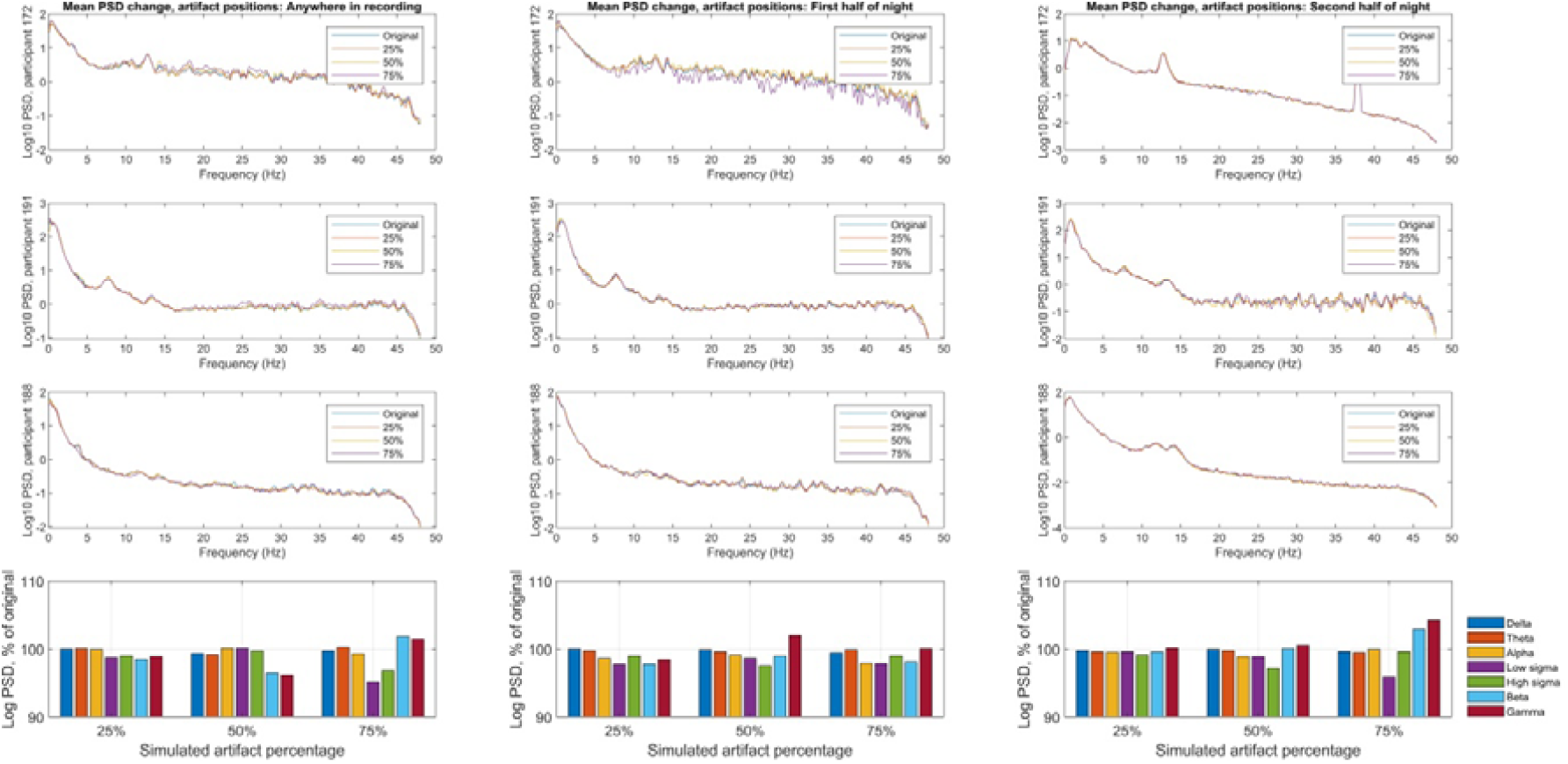
The effect of false-positive artifacts on PSD estimates from the representative channel C3 in NREM. **Panel A:** PSD estimates from three randomly selected participants after randomly setting 0% (original PSD estimate), 25%, 50% or 75% of epochs (separate lines) from either anywhere during the night (left panels), from the first half of the night (middle panels), or from the second half of the night (right panels) to false positive artifacts. **Panel B:** grand average (across all participants) bandwise PSD estimates after the exclusion of false positives, expressed relative to the original.

**Figure 7.**
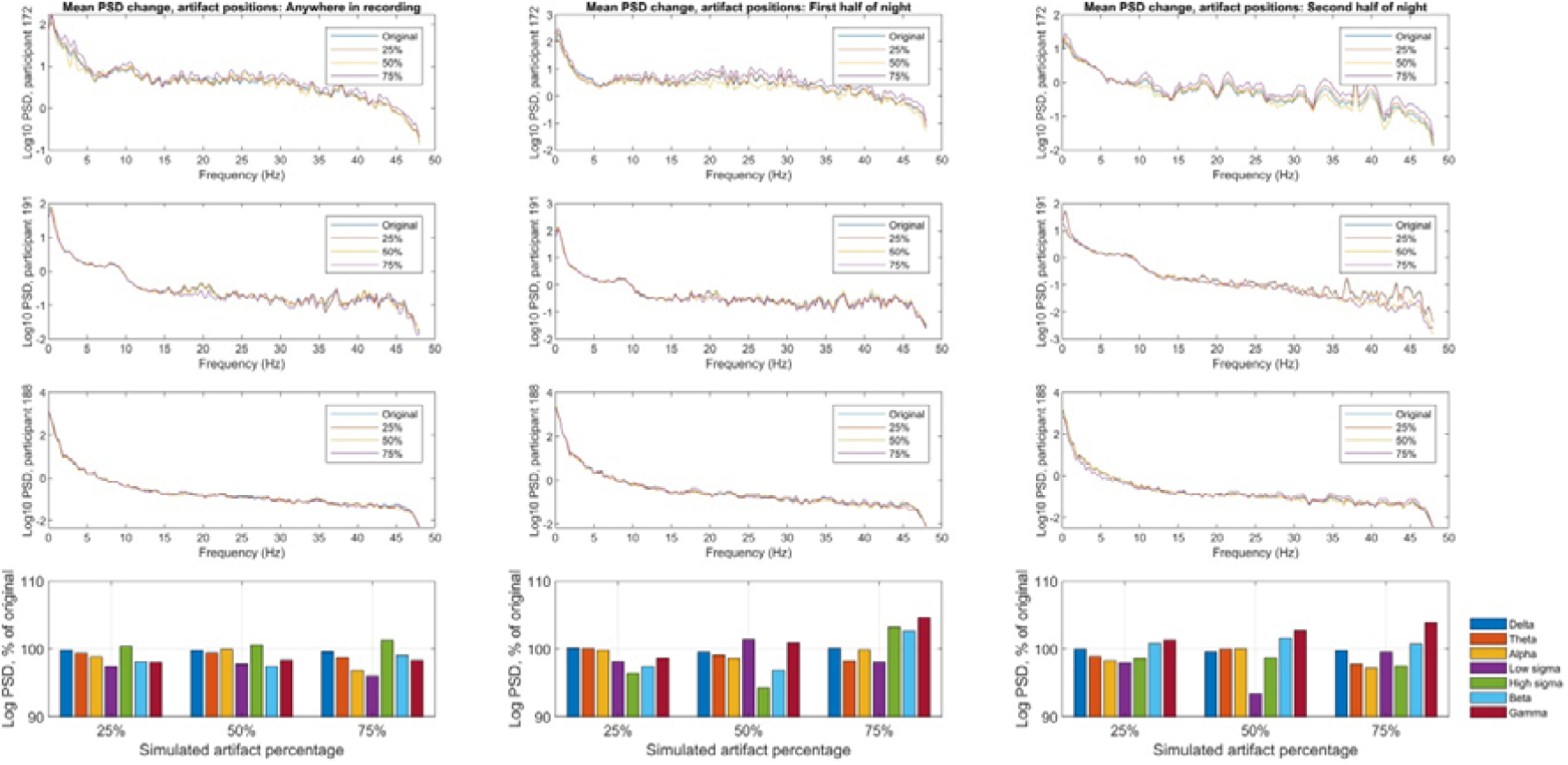
The effect of false-positive artifacts on PSD estimates from the representative channel C3 in REM. **Panel A:** PSD estimates from three randomly selected participants after randomly setting 0% (original PSD estimate), 25%, 50% or 75% of epochs (separate lines) from either anywhere during the night (left panels), from the first half of the night (middle panels), or from the second half of the night (right panels) to false positive artifacts. **Panel B:** grand average (across all participants) bandwise PSD estimates after the exclusion of false positives, expressed relative to the original.

Overall, minimal changes in PSD estimates were observed regardless of the settings used to simulate false positive artifacts. The mean change (across all conditions and frequency bands) was 0.5%, the mean absolute change was 1.16%, with range [-8% - 4.8%], indicating that small subsets of the actually available data epochs provide reliable approximations of the true PSD average even after the majority of data is excluded.

## Discussion

The processing of large-scale sleep EEG databases is only feasible with automatic methods. In our work, we systematically explored 1) the effect of the erroneous inclusion of artefactual data to PSD estimates, 2) the effect of data loss, including time-biased data loss, on PSD estimates, 3) the similarity of visual detections to the results of a simple automatic algorithm previously used in large datasets (Purcell et al., 2017; Djonlagic et al., 2021), 4) the similarity of the shape and external correlations of PSD estimates resulting from the use of visual or automatic detectors.

We found that while visual scorers marked a substantial percentage of epochs (8% in NREM and 18% in REM) as artefactual, the vast majority of these artifacts would have had little or no effect on PSD estimates if left undetected. The peak of the histogram of artifact effects was at zero, and the distribution of artifact effects was symmetrical for most frequency ranges, except for beta and gamma in both vigilance states and for NREM delta. In other words, artifacts tend to have, on average, a neutral effect on PSD estimates, with the exception of high frequencies where muscle artifacts tend to result in overestimates of PSD and NREM delta where artefactual epochs contain less of usual low-frequency activity that characterizes this vigilance state.

However, a substantial minority of artifacts had significant effects on PSD estimates, resulting in a change of several percent due to the erroneous inclusion of a single artefactual epoch. Such artifacts mostly affected the lowest and highest frequency activities (see **Figure 1** and **Figure 2** for an illustration of the high variability of PSD estimates at these frequencies as a result of including artifacts). In alternate analyses looking at the between-participant similarity of PSD estimates with or without artifact detections (**Figure 3** and **Figure 4**), we also found that artifacts mostly affected PSD estimates above the spindle frequency range, likely due to them resulting from high-amplitude muscle artifacts (Brunner et al., 1996; Yuval-Greenberg and Deouell, 2009). This mirrors previous findings (Malafeev et al., 2019), which also reported that high frequencies are the most sensitive to artifacts. Interestingly, PSD estimates at or below the spindle were very well recovered in NREM (r>0.9, **Figure 3**) even if no artifact detection was used at all. External correlations with age and sex (**Figure 5**) were also reasonably well recovered in this frequency range from PSD estimates calculated without considering artifacts.

Our findings about the effect of visually detected artifacts suggest that manual scorers, at least in our sample, are too inclusive and mark up many data epochs as artefactual in which no anomalies are present which would significantly affect PSD estimates. This is especially true for the middle frequency ranges, including the spindle frequency range, where physiological activity likely overpowers all but the most powerful artifacts.

However, low and especially high frequencies are still affected by artifacts, a significant minority of which causes substantial changes in the PSD estimates by themselves. We employed a very simple artifact detection algorithm, using extreme values of the Hjorth parameters (with a set of thresholds) of epochs to identify anomalous data. We found that using this method with any setting resulted in substantial improvements in the quality of PSD estimates. There was no clear threshold setting which resulted in the best performance. The rate of visual artifact detection was best approximated with dual thresholds and intermediate stringency (2.5-2.5 SD). However, this criterion is dubious because, as previously discussed, a large proportion of visually detected artifacts does not make meaningful differences in PSD estimates. Agreement with visual detections was lowest with the most stringent detectors, however, it remained relatively constant with all less stringent methods. All detectors resulted in high (r>0.9, typically r>0.95) correlations between PSD estimates using visual and automatic artifact detections **(Figure 3 and Figure 4)**, highlighting their utility especially in accurately recovering high-frequency PSD estimates. While no clearly superior algorithm emerged, dual-threshold settings had a tendency to especially well approximate visually-scored PSD estimates at high frequencies. External correlations with age and sex **(Figure 5)** were also well recovered with any automatic detection method, although the most extreme (and possibly most accurate) effects were typically found with visual detections.

Our work has a number of limitations. Most significantly, we investigated full-night laboratory PSG recordings of reasonably high quality. The effect of artifacts and the difficulty of identifying them is possibly different in other types of recordings, for example, in routine wake EEG, during naps, or in recordings performed with mobile EEG headbands (Arnal et al., 2020). Another limitation is the use of PSD estimates as the benchmark of artifact detection quality. We chose PSD because it is a frequently used simple EEG marker with well-known correlates most researchers are familiar with. A large number of other metrics (Stancin et al., 2021), such as time-frequency coupling, coherence, entropy or fractal dimensions can be calculated from EEG, and specific waveforms such as slow waves and spindles can be detected, and other metrics may be differently affected by artifacts.

A global evaluation of our findings suggests that artifacts in night sleep EEG recordings do not represent a critical problem. Most artifacts minimally affect PSD estimates (and vice versa, PSD estimates are accurately recovered if only less than a fourth of all data is available for calculations, even if artifacts are heavily concentrated in only one half of the night). PSD is estimated with reasonable accuracy up to the spindle frequency range even without any regard for excluding artifacts. The remaining problems resulting from artifacts are mostly eliminated with the use of a simple algorithm, with no substantial difference resulting from algorithm settings. These findings are encouraging for the prospects of large, freely available datasets. In these datasets, the meticulous visual observation of data is not feasible, access to sophisticated detection algorithms may be limited to some researchers, and the proliferation of detection algorithms may cause issues with reproducibility. Our findings – in line with previous findings exploring other algorithms (t Wallant et al., 2016; Mariani et al., 2018; Malafeev et al., 2019; Saifutdinova et al., 2019) – suggest that accurate PSD estimates strongly resembling the output of visual scoring can be achieved by very simple detection algorithms, and with some limitation even without any regard from artifact detection.

Limitations of the study

## Acknowledgments

This work was supported by the Ministry of Innovation and Technology of Hungary from the National Research, Development and Innovation Fund, financed under the TKP2021-EGA-25 funding scheme and by the grant OTKA PD 138935.

## Declaration of interests

All authors declare no conflict of interest.

## Notes

https://zenodo.org/record/7934586

### Competing Interest Statement

The authors have declared no competing interest.

