## Supplementary figure for "Comparing manual and automatic artifact detection in sleep EEG recordings"

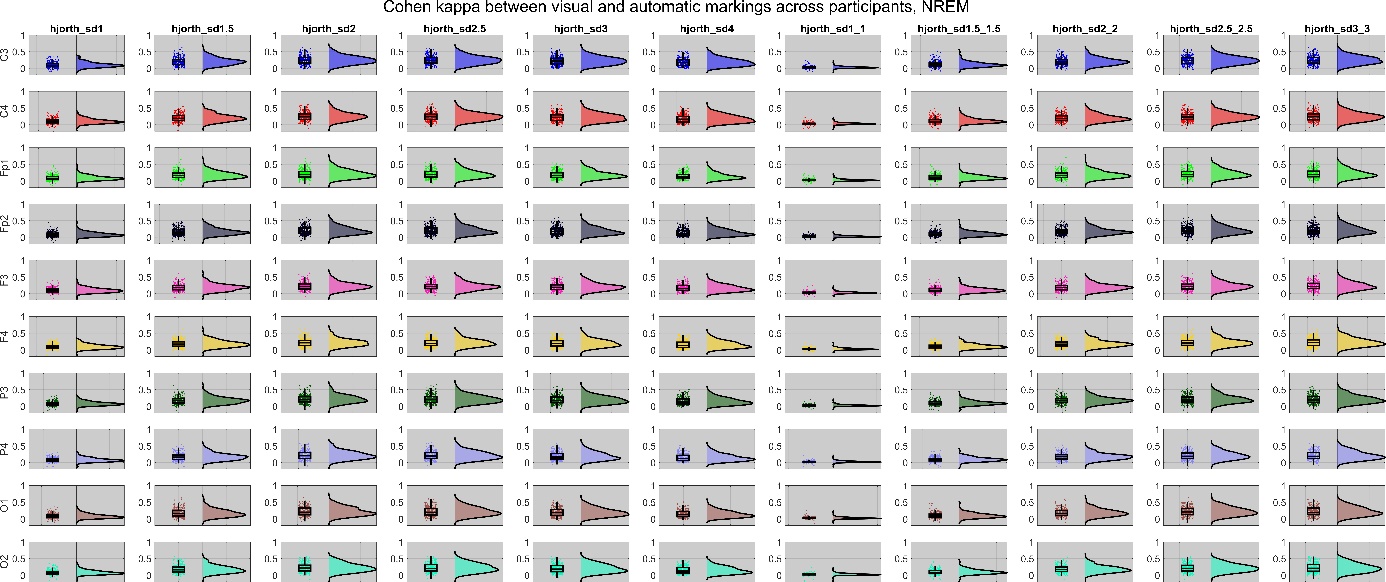

**Supplementary Figure S1**. The similarity of visual and automatic artifact detections in NREM. Similarity is expressed as Cohen’s kappa between scorings of visual and automatic detectors. Each panel shows similarity using a different Hjorth parameter z-score threshold (columns) on a different channel (rows). The left side of each panel shows the correlations in each participant using data points overlain with a box plot. The right side is a kernel density histogram illustrating the distribution of correlations across participants.

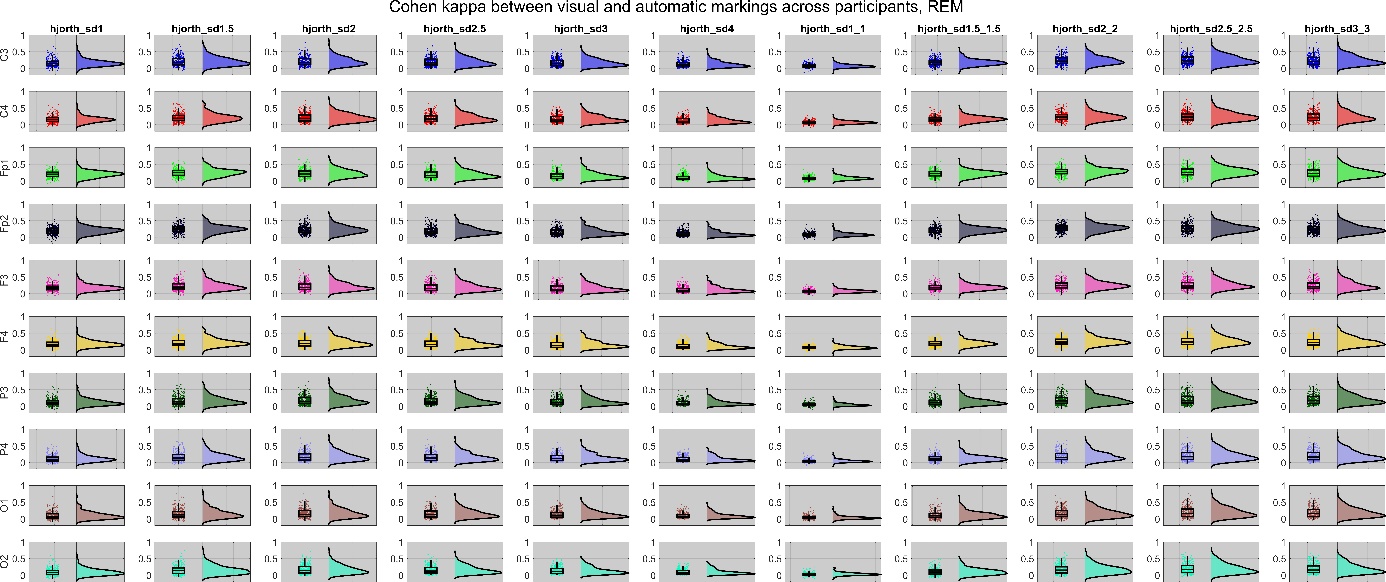

**Supplementary Figure S2**. The similarity of visual and automatic artifact detections in REM. Similarity is expressed as Cohen’s kappa between scorings of visual and automatic detectors. Each panel shows similarity using a different Hjorth parameter z-score threshold (columns) on a different channel (rows). The left side of each panel shows the correlations in each participant using data points overlain with a box plot. The right side is a kernel density histogram illustrating the distribution of correlations across participants.
